# *Aedes albopictus* colonies from different geographic origins differ in their sleep and activity levels but not in the time of peak activity

**DOI:** 10.1101/2024.03.15.585187

**Authors:** Nicole E. Wynne, Emilie Applebach, Karthikeyan Chandrasegaran, Oluwaseun M. Ajayi, Souvik Chakraborty, Mariangela Bonizzoni, Chloé Lahondère, Joshua B. Benoit, Clément Vinauger

**Affiliations:** Department of Biochemistry, Virginia Polytechnic Institute and State University, Blacksburg, VA 24061, USA; Department of Biological Sciences, University of Cincinnati, Cincinnati, OH 45221, USA; Department of Biology and Biotechnology, University of Pavia, Pavia 27100, Italy

**Keywords:** Disease vector insects, Mosquitoes, Activity patterns, Sleep patterns, *Aedes albopictus*, Intra-specific Variability

## Abstract

Mosquitoes occupy a wide range of habitats where they experience various environmental conditions. The ability of some species, such as the tiger mosquito, *Aedes albopictus*, to adapt to local conditions certainly contributes to their invasive success. Among traits that remain to be examined, mosquitoes’ ability to time their activity with that of the local host population has been suggested to be of significant epidemiological importance. However, whether different populations display heritable differences in their chronotype has not been examined. Here, we compared laboratory strains originating from 8 populations from 3 continents, monitored their spontaneous locomotor activity patterns, and analyzed their sleep-like states. Overall, all strains showed conserved diurnal activity concentrated in the hours preceding the crepuscule. Similarly, they all showed increased sleep levels during the morning and night hours. However, we observed strain-specific differences in the activity levels at each phase of the day. We also observed differences in the fraction of time that each strain spends in a sleep-like state, explained by variations in the sleep architecture across strains. Human population density and the latitude of the site of geographic origin of the tested strain showed significant effects on sleep and activity patterns. Altogether, these results suggest that *Ae. albopictus* mosquitoes adapt to local environmental conditions via heritable adaptations of their chronotype.

## Introduction

Mosquitoes are present on all but one continent, imposing a significant epidemiological burden on human populations by transmitting pathogens causing human diseases. *Aedes aegypti* and *Aedes albopictus* are the primary vectors of epidemiologically relevant viruses, such as the yellow fever virus, dengue virus, chikungunya virus, and Zika virus (Lounibos and Kramer 2016). These two *Aedes spp.* species are anthropophilic, meaning they prefer humans over other hosts, and blood-feed in multiple small meals from different hosts, which increases their chances of transmitting arboviruses (Kamal et al. 2018). *Aedes spp*. mosquitoes are also highly invasive and have spread out of their native range, Africa for *Ae. aegypti* and South East Asia for *Ae. albopictus,* to now be globally distributed. While *Ae. aegypti* is mostly a tropical and subtropical species, *Ae. albopictus* has established populations in both tropical and temperate areas of the world (Lounibos and Kramer 2016, Manni et al. 2017, Crawford et al. 2023). With the effects of climate change, it is expected that regions suitable for *Ae. albopictus* will increase by 50% by the end of the century, exacerbating the risk of mosquito-borne diseases worldwide (Rochlin et al. 2013). In this context, there is a clear need for an improved understanding of the biological traits contributing to *Aedes spp*. ability to adapt to such a range of environmental conditions.

Amongst plastic traits, the ability of mosquitoes to adapt to different day lengths, temperatures, and climates has been extensively discussed as an important contributor to their invasive success (Rund et al. 2012, Reinhold et al. 2018, Upshur et al. 2019, Benoit and Vinauger 2022). Using various methods of quantification, including field collections and direct human landing observations, multiple studies revealed differences between the activity patterns of *Ae. albopictus* populations, although chronobiology has received little attention compared to other aspects of mosquito biology. For instance, *Ae. albopictus* from Mercer County in New Jersey (USA) and Volusia County in Florida, (USA) were shown by collection through human sweep netting and/or carbon dioxide-baited rotator traps to display unimodal activity patterns peaking during solar noon (Unlu et al. 2021). Overall, the suburban population from Florida exhibited lower activity levels than the urban population from New Jersey. Using a two-layer mosquito net with human bait, Delatte *et al*. compared *Ae. albopictus* activity inside and outside of houses on the tropical French island of La Réunion (Delatte et al. 2010). Both sub-groups had bimodal activity patterns peaking right after sunrise and right before sunset. Higa et al. collected *Ae. albopictus* from multiple sites around Nagasaki (Japan) during four phases of the day (Higa et al. 2000). While mosquitoes displayed a bimodal activity pattern with a peak before sunrise and one before sunset, these Japanese populations also displayed increased activity during the night. This nocturnal activity has also been reported in laboratory experiments, particularly in host-seeking females (Yee and Foster 1992, Barnard et al. 2011).

Besides spontaneous locomotor activity, patterns of sleep-like states are another important component of mosquitoes’ chronotype because sleep deprivation is correlated with reduced host landing and blood-feeding propensity in *Ae. aegypti* (Ajayi et al. 2022). Sleep-like states in *Aedes*, *Culex*, and *Anopheles* mosquitoes are characterized by stereotypical postural changes, increased arousal thresholds, prolonged immobility during specific periods of the day, and sleep rebound after prior deprivation (Ajayi et al. 2020, 2022, 2023). In the fruit fly, *Drosophila melanogaster,* even though sleep is a highly conserved physiological function, extensive variations in sleep duration or pattern have been observed both among species and between individuals of the same species (Harbison et al. 2013). Transcriptomic studies and artificial selection experiments identified the genetic networks that underlie natural variations in sleep within populations (Svetec et al. 2015, Harbison et al. 2017, Serrano Negron et al. 2018). However, before the molecular basis of sleep variation can be unraveled in mosquitoes, their phenotype-level variation in sleep remains to be quantified in a standardized manner.

Understanding whether chronotypes differ across mosquito populations is important to develop efficient control strategies. For example, previous studies on *Ae. aegypti*, highlighted the importance of applying adulticide treatments at the proper time of day based on the targeted mosquito population’s activity (Wilke et al. 2023). In the malaria vector *Anopheles gambiae,* strain-specific differences in total daily activity were also observed (Rund et al. 2012). However, discrepancies between studies are preventing direct comparisons between populations, especially within the same species. Indeed, mosquito chronobiology can be quantified using different approaches, such as field captures, direct observations by the experimenter, and various laboratory assays, which vary in their sensitivity and temporal resolution; thus, results are not directly comparable.

As a first step towards an improved understanding of the intraspecific variability in mosquito chronobiology, we present here a controlled, laboratory-based comparative analysis of the locomotor activity of multiple *Ae. aegypti* (see companion paper, Ajayi et al. 2024) and *Ae. albopictus* (the present study) strains originating from different parts of the world. Using standardized activity monitoring assays, we tested whether adaptations to the particular environmental conditions of origin are reflected in mosquitoes’ spontaneous locomotor activity and sleep patterns. Taking advantage of the fact that our eight *Ae. albopictus* strains had been acclimated to laboratory colony conditions, these experiments also allowed us to test whether habitat-driven differences in chronotypes would persist under laboratory conditions, *i.e.,* whether they are heritable. Our results show that diurnal behavior, with a peak in activity at the end of the day, is highly conserved in this species. Still, significant differences in activity levels during various phases of the day and in the number of sleep bouts were observed across strains. Our analysis suggests that human population density, more than the latitude of the location of origin, significantly contributes to explaining the observed differences.

## Materials and Methods

### Mosquitoes

Eight *Ae. albopictus* strains were used throughout the experiments: Blacksburg (Virginia, USA - established in 2021), ATMNJ95 (Keyport, New Jersey, USA, NR-48979, ATCC®, Manassas, VA, USA, established in 1995), Gainesville (Florida, USA, MRA-804, ATCC®, Manassas, VA, USA), Jacksonville (Florida, USA), Brunswick (Georgia, USA), Crema (Italy - established in 2017 (Carlassara et al. 2023)), Foshan (China - established in the 1980s (Carlassara et al. 2023)), and Kyushu (Japan).

The Blacksburg, Crema, Foshan, and Gainesville colonies were maintained at Virginia Tech in climatic chambers set at 25 ± 1 °C, 60 ± 10% relative humidity (RH), and under a 16-8 h light–dark (L/D) cycle. Cages of adults were fed weekly using an artificial feeder (D.E. Lillie Glassblowers, Atlanta, Georgia; 2.5 cm internal diameter) with heparinized bovine blood (Lampire Biological Laboratories, Pipersville, PA, USA) heated at 37 °C using a circulating water-bath. Between blood-meals, mosquitoes were fed *ad libitum* with 10% sucrose. No vertebrate hosts were used to rear mosquitoes, nor during the experiments. Eggs were collected from blood-fed females and hatched in deionized water. Larvae were reared in groups of 200 in covered pans (26 × 35x4 cm) containing deionized water and fed daily with fish food (Hikari Tropic 382 First Bites—Petco, San Diego, CA, USA). Pupae, in groups of 100, were isolated in 16 oz containers (Mosquito Breeder Jar, Bioquip Products, Rancho Dominguez, CA, USA) until emergence. For all experiments, pupae were collected and placed into chambers with a 12-12 h light-dark (L/D) cycle. Then, mated female mosquitoes from these cages were used for experiments when they reached 3-5 days old (*i.e.*, when they reached host-seeking age (Tallon et al. 2019)).

The Jacksonville, Brunswick, Kyushu, Keyport, and Gainesville (which served as a control for the testing site) colonies were maintained at the University of Cincinnati at 25°C, 80% relative humidity (RH) under a 15 h:9 h light: dark (L/D) cycle with access to water, and 10% sucrose *ad libitum*. Eggs were produced from 2- to 4-week-old females through blood feeding with a human host (University of Cincinnati IRB 2021-0971). After egg hatching, larvae were separated into 18×25×5 cm containers (at a density of 250 individuals per container). They were fed finely ground fish food (Tetramin, Melle, Germany) and maintained in an incubator at 24°C, 70–75% RH, under a 12 h:12 h L/D cycle until adult emergence. Emerged adult mosquitoes had access to water and 10% sucrose *ad libitum*. All adult female mosquitoes used for the experiment were 7–14 days post-ecdysis. As the experimenters represent potential hosts to the mosquitoes, the experiment was conducted in an enclosed room in an isolated building that was not accessed during experiments to eliminate potential disturbances.

The Gainesville strain was maintained and tested at both universities to control for the potential effects of differences in rearing conditions between the two laboratories (see below for details). All experiments were conducted for a minimum of 6 consecutive days, and the first day was discarded from the analysis to control for potential confounding effects of the handling of mosquitoes.

### Activity assays

Mosquito activity was measured using activity monitors that consist of 32 openings, with three sets of infrared emitters and detectors per opening (Trikinetics LAM25, Waltham, MA, USA.). Each mosquito was placed into a clear cylindrical glass tube with mesh covering one end and a custom-made cotton plug with access to 10% sucrose solution on the other end (See (Eilerts et al. 2018, Upshur et al. 2019, Ajayi et al. 2022) for details). These tubes were then placed into the openings of the monitor so that the infrared beams bisected each tube in the middle. The whole activity monitor was then placed into a temperature-controlled chamber (model I-36VL, Percival Scientific, Perry, Iowa, USA), maintaining a temperature of 25 ± 1 °C and 40 ± 10% relative humidity (RH) throughout the experiments. Mosquitoes were maintained under a 12-12 h light-dark cycle with a 1 h long soft transition between light and dark to mimic sunrise (ZT 23:30-00:30) and sunset (ZT 11:30-12:30). Once set up, the mosquitoes were left undisturbed in these tubes for ten days. Daily activity was recorded as the number of beam crossings every minute using the DAMSystem3 Software (Trikinetics, Waltham, MA, USA). At the end of the ten days, the monitors were removed from the chamber, and mosquitoes were examined for survival. The data from tubes containing mosquitoes found dead at the end of the experiment (30 ± 10% across experiments) were discarded from the analysis. To remove potential outliers, the 5% of individuals with the least and highest number of beam breaks were excluded from the analysis.

### Statistical Analysis

In actometer experiments, each mosquito represents an independent replicate. We performed actometer assays to achieve n > 45 mosquitoes for each strain, and this sample size is comparable to similar assays in previous studies (Eilerts et al. 2018, Ajayi et al. 2020, 2022). Detailed sample sizes for each strain are provided in the results section. Beam count data from the actometer was imported in R (version 4.3.1) and used to compute the activity level, sleep duration, and number of sleep bouts either as a function of the time of day or binned into 4 phases of equal duration (*i.e.*, 6 hours): sunrise (ZT 22 - 4), day (ZT 4-10), sunset (ZT 10-16), and night (ZT 16-22). These response variables were obtained using built-in functions of the *Rethomics* package (Geissmann et al. 2019).

Differences in activity and sleep levels between strains and phases of the day were evaluated using ANOVA followed by Tukey’s post-hoc tests for multiple comparisons of means with a Bonferroni adjustment of the *p*-values. The effect of the predictor variables such as the strain identity, the latitude, and the population density of the location of origin (obtained from www.census.gov for US cities and www.citypopulation.de for Italy, China, and Japan) was evaluated using generalized linear models computed with the *glm* function of the *lme4* package (Bates et al. 2014). Comparisons between least-square means of the response variables across the predictor variables were performed using the *emmeans* package (Lenth et al. 2019).

## Results

### Testing the same strain at different locations shows comparable activity profiles

We selected 8 strains of laboratory-reared mosquitoes originating from populations located in Asia, Europe, and North America to evaluate intra-species differences in chronobiology (**Figure 1a-b**). The selected strains originate from a range of latitudes (from 23.02°N to 45.36° N) where different population densities are observed (from 2060.23 to 16575.92 inhabitants per square km, **Figure 1c**); latitude and population density are not correlated (Pearson’s product-moment correlation, *t* = -0.29269, df = 7, *p*-value = 0.7782). A total of 524 females were used for these experiments (Blacksburg: n = 53, Brunswick: n = 45, Crema: n = 96, Foshan: n = 85, Gainesville: n = 90, Jacksonville: n = 45, Keyport: n = 56, Kyushu: n = 54).

**Figure 1.**
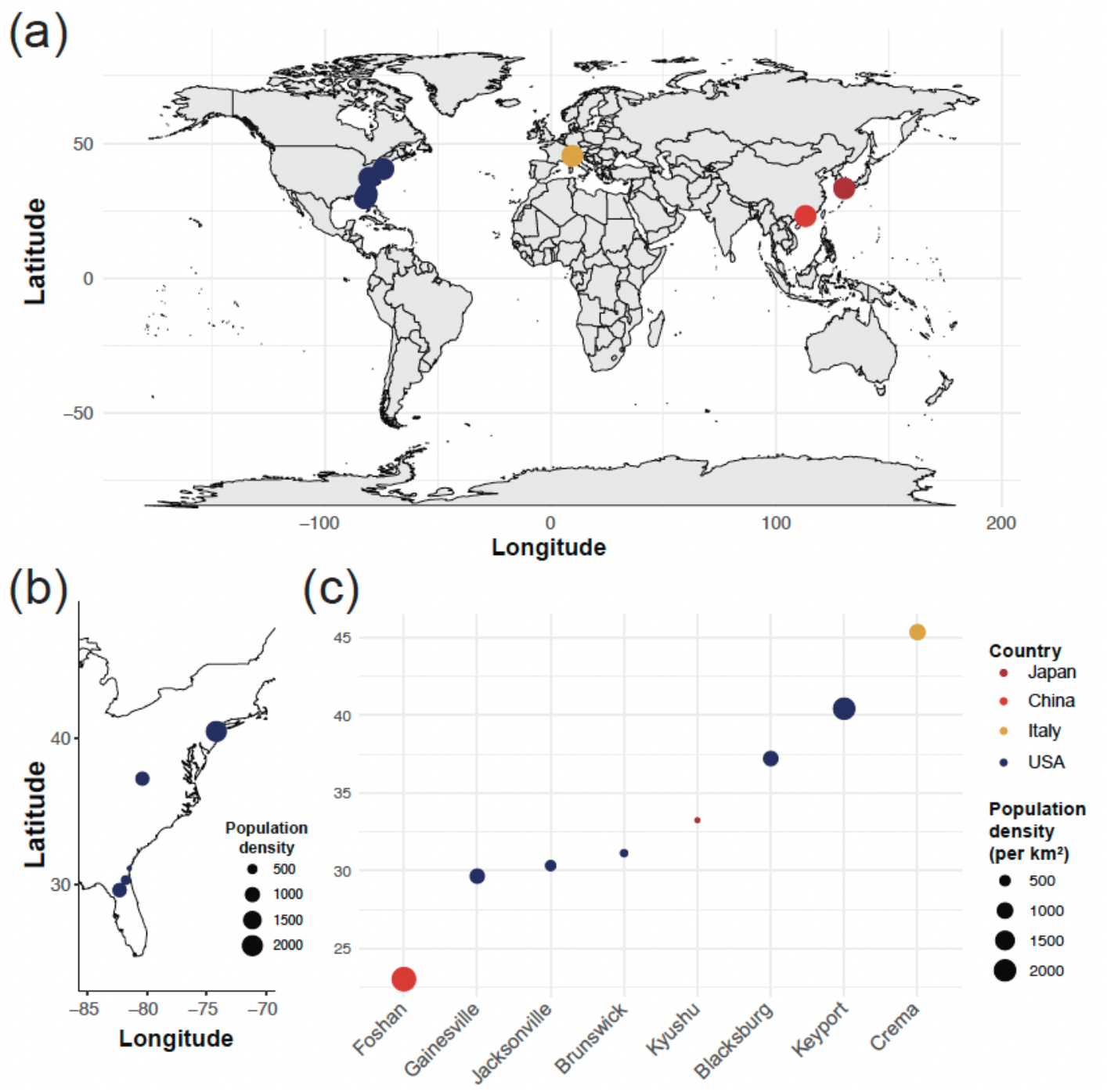
Distribution and host populations’ characteristics of the tested strains. **(a)** World map depicting the geographic origin of the 8 *Aedes albopictus* strains tested in the study. **(b)** Detailed location of the US populations’ geographic origins. The size of the point encodes for the population density at the collection site (in inhabitants per square kilometer). **(c)** Latitude of the collection site for each of the 8 strains, color-coded by country of origin and size-coded for population density (in inhabitants per square kilometer).

To ensure that experimental results were not biased by the testing site (*i.e.*, Virginia Tech *vs* the University of Cincinnati), the Gainesville strain was studied at both locations. Experiments conducted on Gainesville mosquitoes at the University of Cincinnati tended to yield a higher number of beam crossings, suggesting higher activity levels, than the experiments conducted at Virginia Tech, in particular at the transition between the light and dark phase of the experiment (Two sample *t* test, *t* = -0.86935, df = 88, *p* value = 0.387), but differences were not statistically significant. Additionally, the overall waveform was qualitatively similar, with a start in increased activity levels around midday, a more prominent peak at the light/dark transition, low nocturnal activity, and a minor activity peak starting in the hour before the dark/light transition (**Figure 2a**). We divided the observed number of beam crosses per unit of time by the average number of beam crosses throughout the experiments to normalize for differences in activity levels. We hypothesize that the observed, statistically not significant, differences in activity levels result from slight differences in temperature and humidity conditions in the experimental rooms, from slight differences in the light intensity in particular during the transitions between phases (night/day), or to the differences in the age of the mosquitoes tested at each site. After normalization, the differences between the day-time activity of the two experiments remained not significant (*t*-test, t = -0.9156, df = 86.553, p-value = 0.3624), but a significant difference was observed when comparing activity levels during the light/dark transition specifically (*t*-test, *t* = -6.3801, df = 64.492, *p*-value < 0.001) (**Figure 2b**). However, because the overall ratio between the day and night activities was not significantly different between the two experimental groups (*t*-test, *t* = 1.7897, df = 88, *p*-value = 0.076, **Figure 2c**), we combined them for the remainder of the analysis. All subsequent comparisons were performed on data normalized for the mean activity over the course of the experiments.

**Figure 2.**
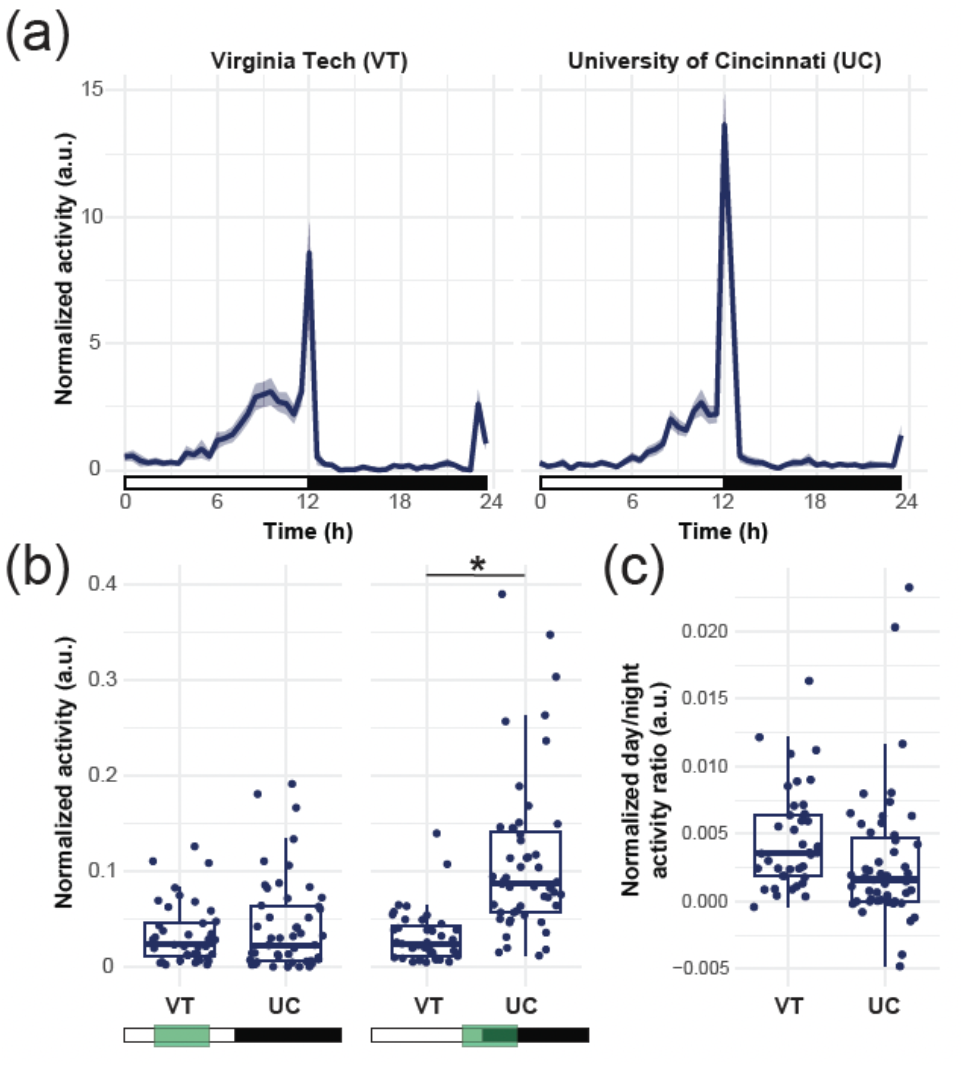
The activity profile of the Gainesville strain is conserved across testing sites, but responses to changes in light sensitivity vary. **(a)** Actograms of the Gainesville strain tested either at Virginia Tech (left) or at the University of Cincinnati (right), showing the normalized average activity profile across the 5 days of the experiment. The white rectangles below the chart indicate the *Zeitgeber* times when the light is ON, and the black rectangles indicate times when the light is OFF. The shaded area around the curve represents the 95% confidence interval. **(b)** Normalized average activity of individual mosquitoes during the day (ZT 4-10, left) or the sunset (ZT 10-16, right). Each point represents an individual mosquito; box plots represent the median, quartiles, and 95% confidence intervals. Green boxes indicate the time windows selected for each comparison. **(c)** Normalized ratio between the day and night time activity of individual mosquitoes. Each point represents an individual mosquito; box plots represent the median, quartiles, and 95% confidence intervals.

### All tested strains show diurnal activity but differ in activity levels

All strains displayed more locomotor activity during the day than during the night, regardless of their geographic origin. However, the waveforms of their activity profiles indicate differences in the relative amount of activity and in the shape of the ramp leading to the main activity peak (**Figure 3, Figure S1**), typically starting around ZT6. All strains displayed a sharp peak at the transition between the light and dark phases (sunset), and most displayed a smaller peak at the transition between the dark and light phases (sunrise). This secondary peak was nearly absent in the Brunswick, Jacksonville, and Kyushu lines, whose activity profile could be described as unimodal. Overall, the Jacksonville strain was the most active, and the Blacksburg strain was the least active among the tested strains.

**Figure 3.**
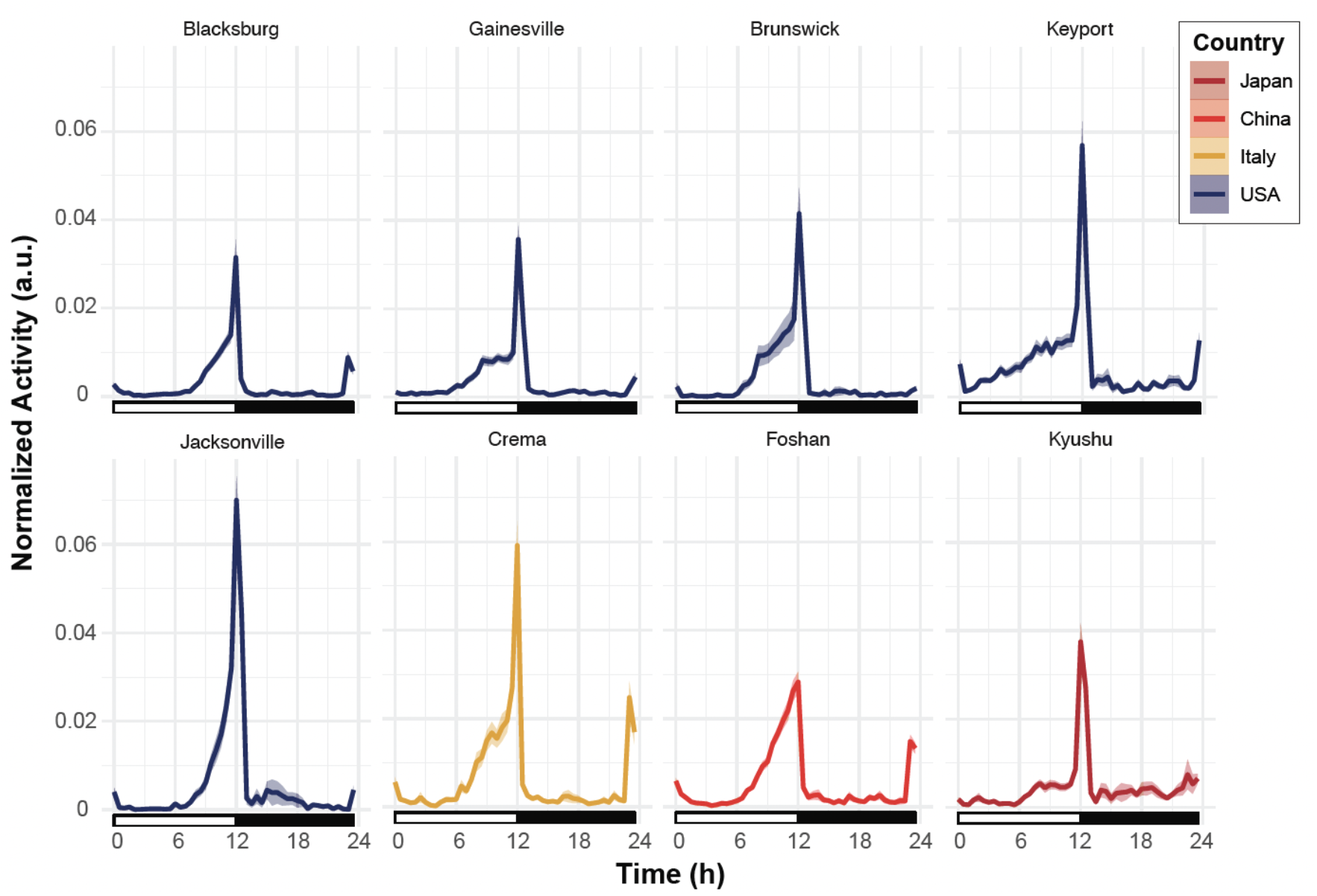
All strains showed diurnal/crepuscular activity. Actograms of the 8 tested strains show the normalized average activity profile across the 5 days of the experiment. The white rectangles below the chart indicate the *Zeitgeber* times when the light is ON, and the black rectangles indicate times when the light is OFF. The shaded area around the curve represents the 95% confidence interval. Each line is color-coded as a function of the country of origin of the population.

A quantitative analysis of the activity throughout the day showed that the strain and the phase were both significant predictors of the activity level (ANOVA, 4 < df < 7, 27.93 < *F* < 278.87, *p* value < 0.001). In accordance with the typical dusk biting activity of the species, all strains were either similarly active at sunset than the least active strain from Blacksburg (Gainesville, Brunswick, Foshan, and Kyushu) or showed significantly higher activity than Blacksburg (Keyport, Jacksonville, Crema) (Tukey post hoc multiple pairwise comparisons, *p* < 0.05). No significant differences were observed among strains’ activity at sunrise and during the night. During the day (*i.e.*, light phase excluding the transition phases), the Keyport, Crema, and Foshan strains were significantly more active than the least active strain, Blacksburg (Tukey post hoc multiple pairwise comparisons, *p* < 0.05). The other four strains were not significantly different from the Blacksburg strain (*p* > 0.05; **Figure 4**).

**Figure 4.**
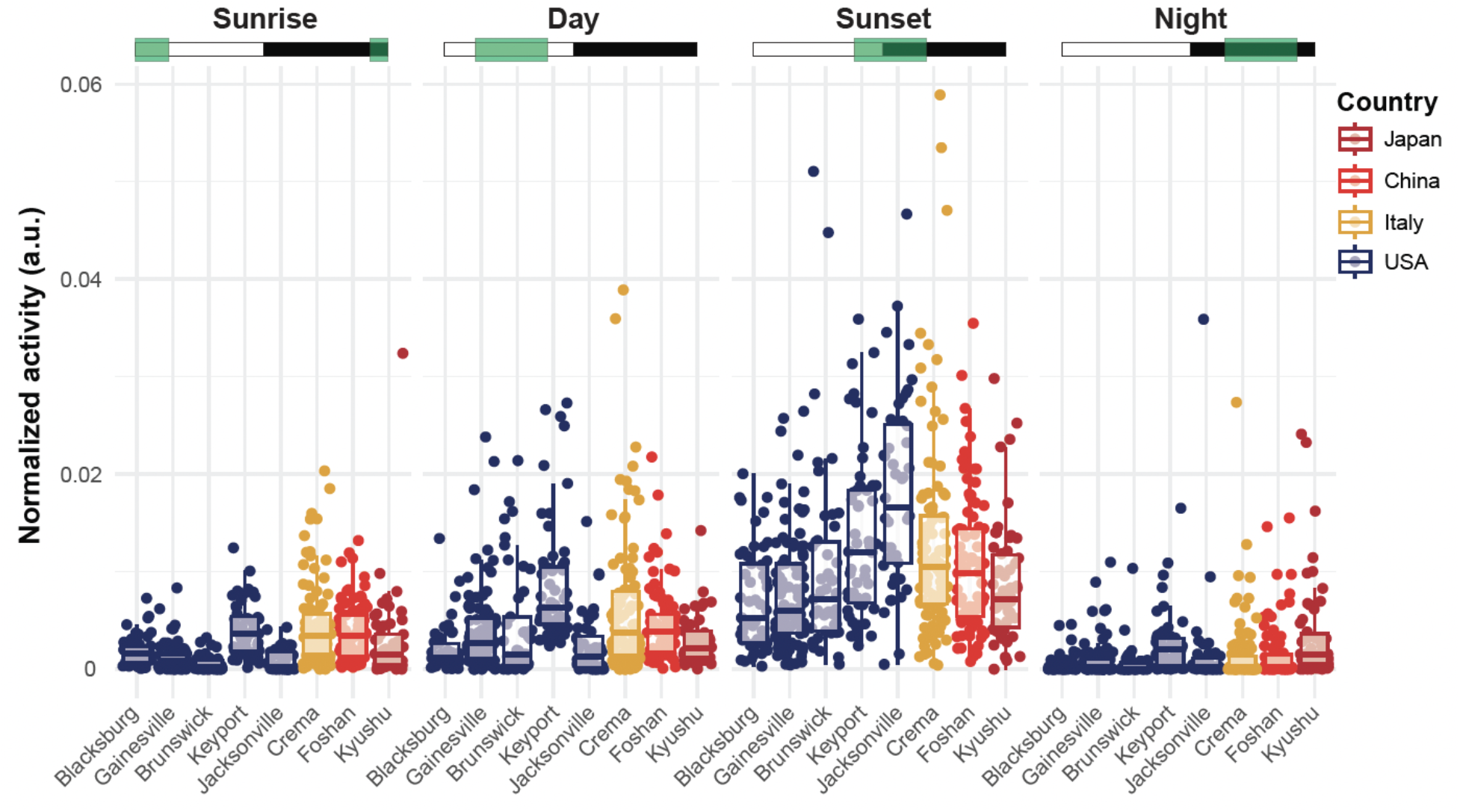
Strains varied in their relative level of activity per phase of the day. Normalized average activity of individual mosquitoes during the sunrise (ZT 22-4), day (ZT 4-10), sunset (ZT 10-16) or night (ZT 16-22) phase. The Green boxes indicate the time windows selected for each comparison subplot, where white rectangles indicate the Zeitgeber times when the light is ON, and the black rectangles indicate times when the light is OFF. Each point represents an individual mosquito; box plots represent the median, quartiles, and 95% confidence intervals. For clarity, significant differences are reported in Supplementary Table 1.

### All tested strains show increased sleep at night but differ in sleep levels during the daytime

Sleep-like states were defined as periods of inactivity (absence of beam crosses) lasting a minimum of 120 minutes (Ajayi et al. 2022). Then, the fraction of time spent asleep throughout the day was calculated for each strain. Similarly to locomotor activity, all strains showed increased sleep levels during the early morning and during the night (**Figure 5**). This pattern was consistent throughout the duration of the experiment (see **Figure S2**), although the shape of the trough in sleep amount around sunset, as well as the overall magnitude of sleep levels, varied across strains.

**Figure 5.**
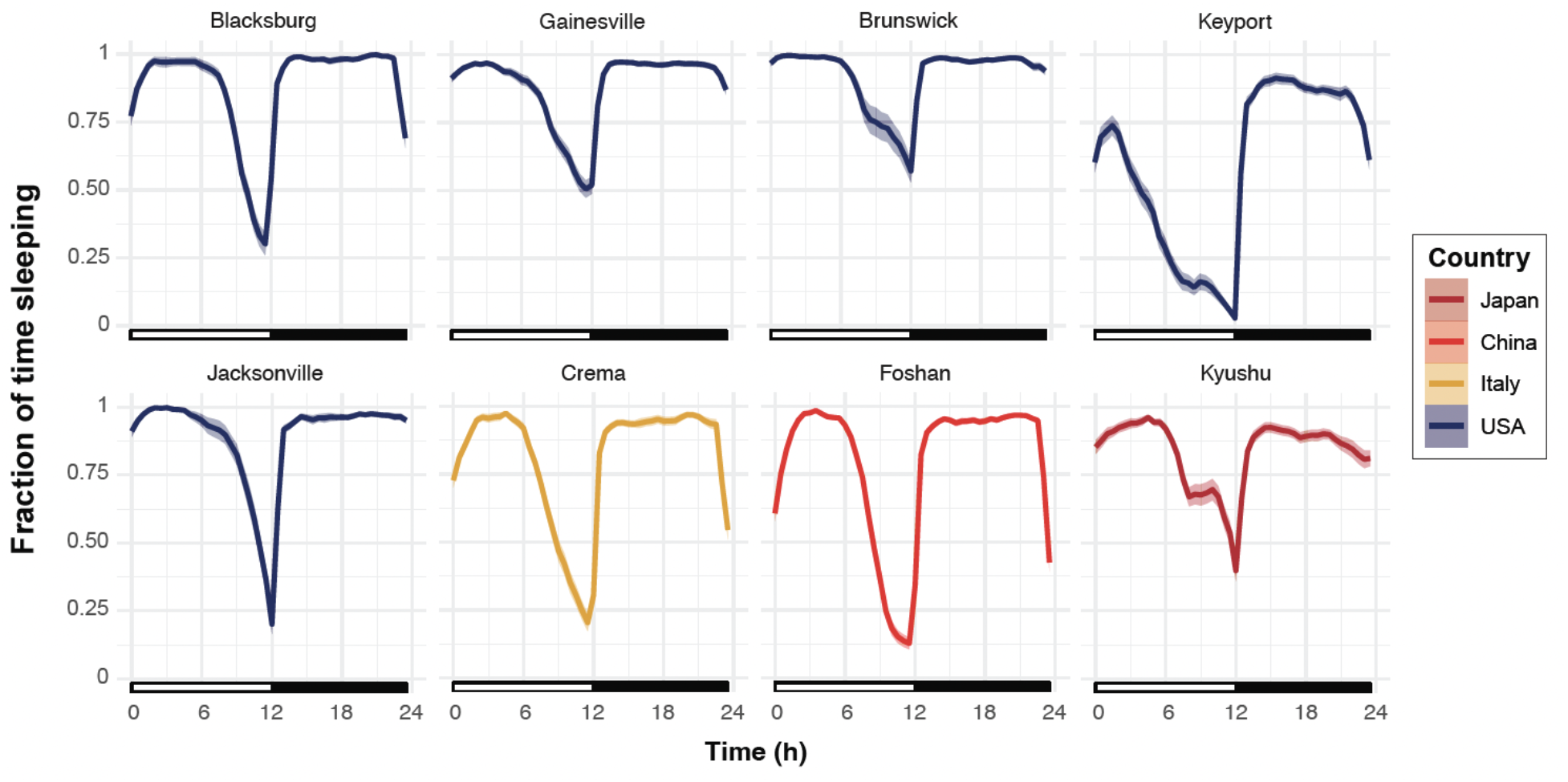
All strains slept more during the morning and night time. Normalized average fraction of time spent in a sleep-like state during the day. The white rectangles below the chart indicate the *Zeitgeber* times when the light is ON, and the black rectangles indicate times when the light is OFF. The shaded area around the curve represents the 95% confidence interval. Each line is color-coded as a function of the country of origin of the population.

Overall, the phase of the day and the strain identity were significant predictors of the fraction of time spent asleep (ANOVA, 3 < df < 7, 144.16 < *F* < 395.45, *p* < 0.001). During the sunrise phase, no significant differences were observed when strains were compared to the least active strain from Blacksburg, except for the Keyport strain, which spent significantly less time asleep (Tukey post hoc multiple pairwise comparisons, *p* < 0.001). However, Crema and Foshan spent significantly less time asleep than the Jacksonville and Brunswick strains, which spent the most time asleep (Tukey post hoc multiple pairwise comparisons, *p* < 0.05) (**Figure 5**). During the day, Foshan, Keyport, and Crema slept significantly less than the Blacksburg strain while the other strains (*i.e.,* Gainesville, Brunswick, Jacksonville, and Kyushi) did not differ significantly from the Blacksburg strain.

Around sunset, all strains slept significantly less than at night, and the Keyport strain slept significantly less than all other strains (Tukey post hoc multiple pairwise comparisons, *p* < 0.05). At night, the Keyport and Kyushu strains spent significantly less time asleep than the Blacksburg strain (Tukey post hoc multiple pairwise comparisons, *p* < 0.05).

### Sleep architecture is conserved in most strains

Differences in the fraction of time spent asleep could be attributed to two main characteristics of the sleep architecture: the number of sleep bouts and the duration of each sleep bout. Overall, the strain identity was a significant predictor of the average sleep bout duration (ANOVA, df = 7, *F* = 12.89, p < 0.001) and of the number of sleep bouts (df = 7, *F* = 92.77, *p* > 0.001) (**Figure 7, Supplementary Table 3,4**). Interestingly, over the course of a day, the Keyport strain showed a significantly higher number of sleep bouts (Tukey post hoc multiple pairwise comparisons, *p* < 0.01) but with significantly shorter bout durations (Tukey post hoc multiple pairwise comparisons, *p* < 0.05) than the Brunswick and Jacksonville strains, which spent a significantly higher fraction of time asleep (**Figure 6**). Finally, all strains showed a similar distribution of sleep bout durations during the day, with more prolonged sleep bouts in the early night (**Figure S3**).

**Figure 6.**
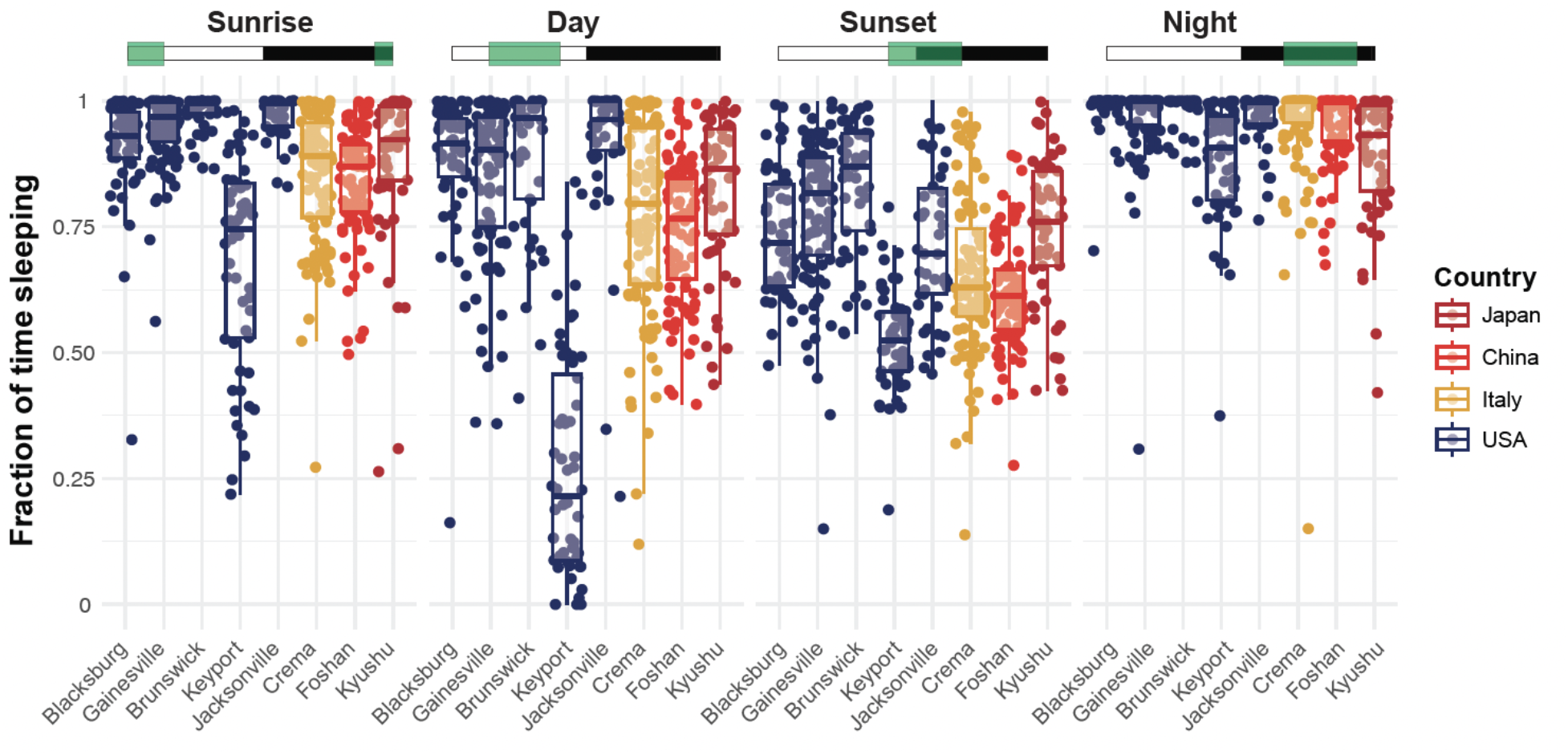
Sleep amounts varied between strains and phases of the day. Average fraction of time spent in a sleep-like state during the day by individual mosquitoes during the sunrise (ZT 22-4), day (ZT 4-10), sunset (ZT 10-16), or night (ZT 16-22) phase. The Green boxes indicate the time windows selected for each comparison subplot, where white rectangles indicate the Zeitgeber times when the light is ON, and the black rectangles indicate times when the light is OFF. Each point represents an individual mosquito; box plots represent the median, quartiles, and 95% confidence intervals. For clarity, significant differences are reported in Supplementary Table 2. Strains are color-coded as a function of the country of origin of the population.

**Figure 7.**
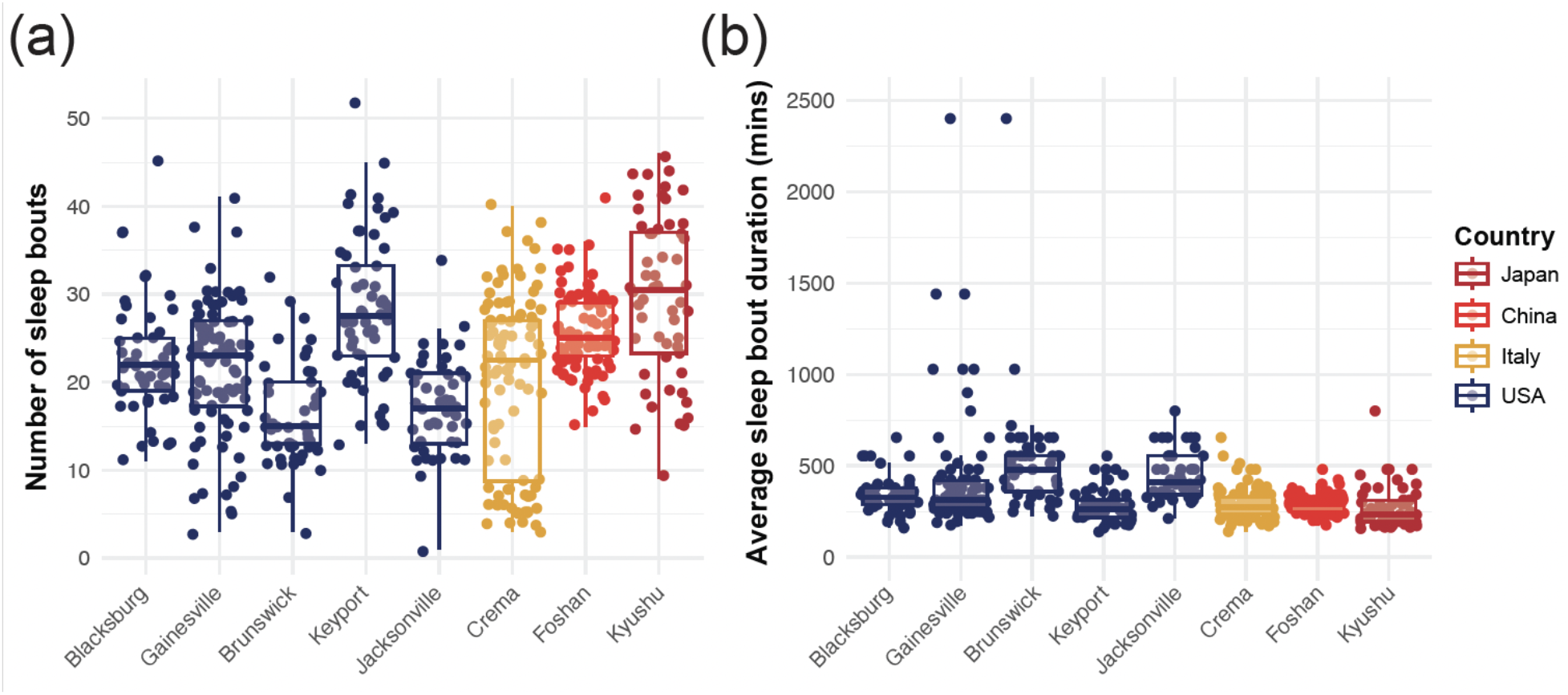
The number of sleep bouts and their average duration are strain-specific. **(a)** Individual mosquitoes’ average number of sleep bouts during each day of the experiment. Each point represents an individual mosquito; box plots represent the median, quartiles, and 95% confidence intervals. **(b)** The average duration of each sleep bout by individual mosquitoes. Each point represents an individual mosquito; box plots represent the median, quartiles, and 95% confidence intervals. Strains are color-coded as a function of the country of origin of the population.

### The geographic origin of the strains explains differences in their activity profile

The latitude and the population density at the geographic origin of the tested strains were independently both significant predictors of the daily activity level (Generalized Linear Model: activity ∼ density + latitude, *p* < 0.001, AIC = -15076), but their interaction was not (Generalized Linear Model: activity ∼ density * latitude, *p* = 0.32, AIC = -15075). The latitude and population densities were significant predictors of the average sleep bout duration, both independently and interactively (Generalized Linear Model: bout duration ∼ density * latitude, *p* < 0.001, AIC = 54185), with the longer bout durations observed at lower population densities and at intermediate latitudes. Interestingly, the latitude, as well as the interactive effect of the latitude and population density, were significant predictors of the number of sleep bouts. Still, the density alone was not a significant predictor of the number of sleep bouts (Generalized Linear Model: number of sleep bouts ∼ density * latitude, *p* < 0.001 for latitude and density: latitude, and *p* = 0.081 for density, AIC = 18589).

## Discussion

The Asian tiger mosquito, *Ae. albopictus,* is native to Asia but has invaded multiple continents with various climates (Bonizzoni et al. 2013, Kotsakiozi et al. 2017, Manni et al. 2017). Previous studies have recorded variations in activity profiles in different populations of *Ae. albopictus.* Many of these studies have used a variety of methods to measure their activity, in particular field-based captures (Delatte et al. 2010, de Lima-Camara 2010, Casas Martínez et al. 2013, Unlu et al. 2021, Wilke et al. 2023). Here, we used activity monitors to record mosquitoes’ activity patterns over a five-day period under constant temperature and humidity conditions, regular light/dark cycles, and in the absence of host cues. This standardized approach limits variations in activity due to the experimental environmental conditions. In addition, all the strains tested here were adapted to laboratory conditions and maintained in a colony for several generations before the experiments. This further controls for potential effects due to differences in larval environments and contributes to isolating effects resulting from selected and inheritable temporal traits.

While testing the same strain, Gainesville, at two different locations revealed some differences in the overall locomotor activity magnitude, the fine-scale architecture of the activity pattern was conserved across sites. Most of the difference resided in the light-to-dark phase transition, where subtle differences in the light ramp used to simulate sunset could explain the observed discrepancies between locations. In addition, the mosquitoes tested at the University of Cincinnati were older than those tested at Virginia Tech. This could potentially contribute to the difference, given that older females may be more inclined to host-seek than their younger counterparts (Tallon et al. 2019, Lau et al. 2020). Of importance, all strains tested at one site (*e.g.*, U. of Cincinnati) did not show higher activity levels than strains tested at the other sites. There is thus a strain specificity in the activity profiles and, likely, in their respective sensitivity to changes in light intensity or age-dependent effects.

All eight strains tested here showed strong activity levels around sunset. However, some of the strains (*e.g.*, Brunswick, Gainesville, Jacksonville, Kyushu) did not show a bimodal activity pattern, with a smaller peak at the end of the night. This confirmed observations made in previous studies where populations from Gainesville, Brunswick, Jacksonville, and Kyushu displayed an unimodal pattern peaking at sunset, similar to the Florida and New Jersey *Ae. albopictus* populations examined in the field, except that these populations displayed a single peak in activity centered around solar noon (Unlu et al. 2021). While significant differences in activity levels were observed throughout the day, most variations were observed during the sunset phase (*i.e.*, starting 2 hours before the transition into the night). In particular, strains originating from Jacksonville and Keyport showed increased activity during this phase compared to other US populations. On the other hand, the Asian and European strains displayed comparably high activity levels at sunset (**Figure 4**).

Similarly, all strains showed high sleep levels during the mid-morning and at night, but not all displayed a strong reduction of sleep levels at sunrise (*e.g.*, Gainesville, Brunswick, Jacksonville; **Figure 5**). In addition, the strain specificity in sleep patterns appears to be decoupled from the strain specificity in activity levels. For example, the Jacksonville strain, which was one of the most active strains, did not show significantly less sleep than the least active US strain from Blacksburg. However, the Keyport strain was the second most active strain and showed the least sleep during the experiment (**Figure 4, 6)**. A deeper analysis of the sleep architecture indicates that this particular strain from New Jersey sleeps in more bouts of shorter duration than its counterpart from Florida (**Figure 7**). This suggests that the genes responsible for regulating activity levels and those coding for characteristics of mosquitoes’ sleep amount and architecture could be part of two independent selective processes.

Although not enough information is available about the exact habitat characteristics of the collection sites used to establish the strains tested here, we used geographic information linked to their populations of origin. Results suggest that the host population and type of environment (by proxy of the latitude of the cities of origin) affect mosquitoes’ sleep and activity profiles. Remarkably, all strains tested here are established laboratory colonies, adapted to feeding on membrane feeders on a similar schedule and maintained under stable laboratory conditions (*e.g.*, temperature, relative humidity) for multiple generations. This indicates that the differences we observed are thus inherited and most likely reflect adaptations to the specific environmental conditions where these mosquitoes were initially captured.

In light of the importance of mosquitoes’ circadian rhythms for synchronizing their sensory performance, physiology, and flight activity with environmental cycles, including patterns of host availability (Gentile et al. 2013, Rund, Bonar, et al. 2013, Rund, Gentile, et al. 2013, Rund et al. 2016, Eilerts et al. 2018, Ajayi et al. 2020), our results suggest that each rhythmic aspect of mosquito biology may be independently selected for. Given the central control of certain insect activities, such as sleep (Shafer and Keene 2021) and the control of olfactory rhythms by peripheral clocks (Jung et al. 2013), this further raises the question of how multiple oscillators coordinate the activity of several behaviors that respond to independent but not disconnected selective forces.

This work sets the stage for future studies (*e.g.*, genomic comparisons, experimental evolution studies) that will be required to understand the mechanisms underlying plasticity and adaptability in mosquito biological rhythms and their contribution to this species’ vectorial and invasive capacities. In particular, direct comparisons of freshly collected mosquitoes with colonized mosquitoes originating from the same population will enable a quantification of the phenotypic and genetic adaptation to laboratory conditions.

Finally, this study shows many similar trends to observation in *Ae. aegypti* based on the accompanying project, where there are minimal lineage-specific effects and differences between experiment locations (Ajayi et al. 2024). The companion paper includes lineages with known differences in ancestry and human preference, and differences in sleep and activity observed for *Ae. aegypti* were shown to vary with other traits for this species.

## Acknowledgements

We thank Helen Oker and Saied “JP” Mirlohi for their help in maintaining the *Aedes albopictus* strains in the colony. The following reagent was obtained through BEI Resources, NIAID, NIH: *Aedes albopictus,* Strain Gainesville, MRA-804, contributed by S. A. Allan; Strain ATM-NJ95, Eggs, NR-48979. Contributed by D. M. Fonseca This study was partially supported by the National Institute of Allergy and Infectious Diseases of the National Institutes of Health under Award Number R01AI155785 (to C.V. for shared incubator space), R01AI148551 (to J.B.B. for shared incubator space), R21AI166633, which focuses on understanding mosquito sleep, and the USDA National Institute of Food and Agriculture, Hatch Research projects VA-1017860 and VA-160212 (to C.V.).

