## Supplementary Information for "*Aedes albopictus* colonies from different geographic origins differ in their sleep and activity levels but not in the time of peak activity"

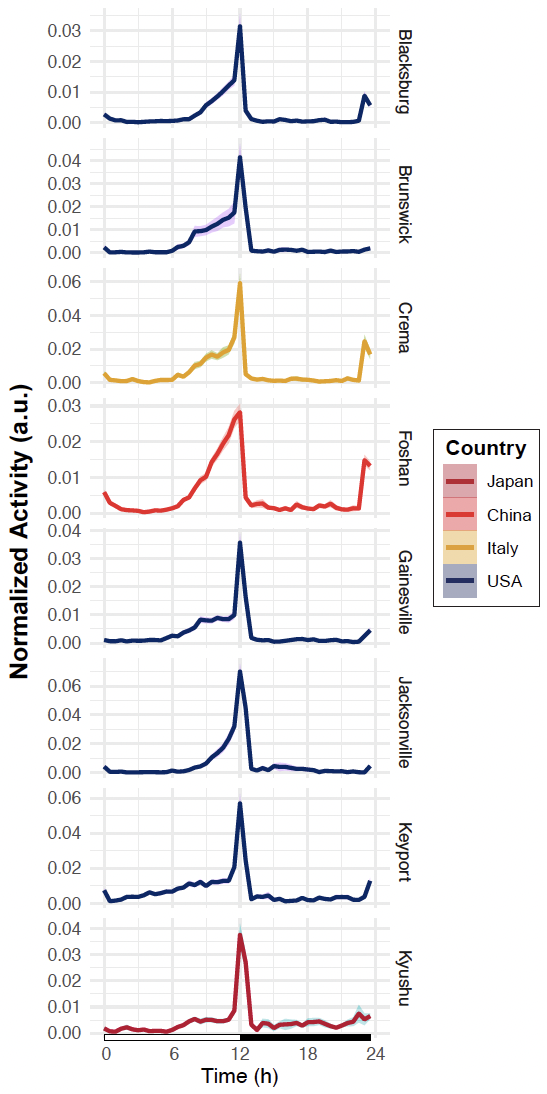

**Figure S1**. Vertically aligned actograms of the 8 tested strains show the normalized average activity profile across the 5 days of the experiment. The white rectangle below the chart indicates the *Zeitgeber* times when the light is ON, and the black rectangle indicates times when the light is OFF. The shaded area around the curve represents the 95% confidence interval. Each line is color-coded as a function of the country of origin of the population.

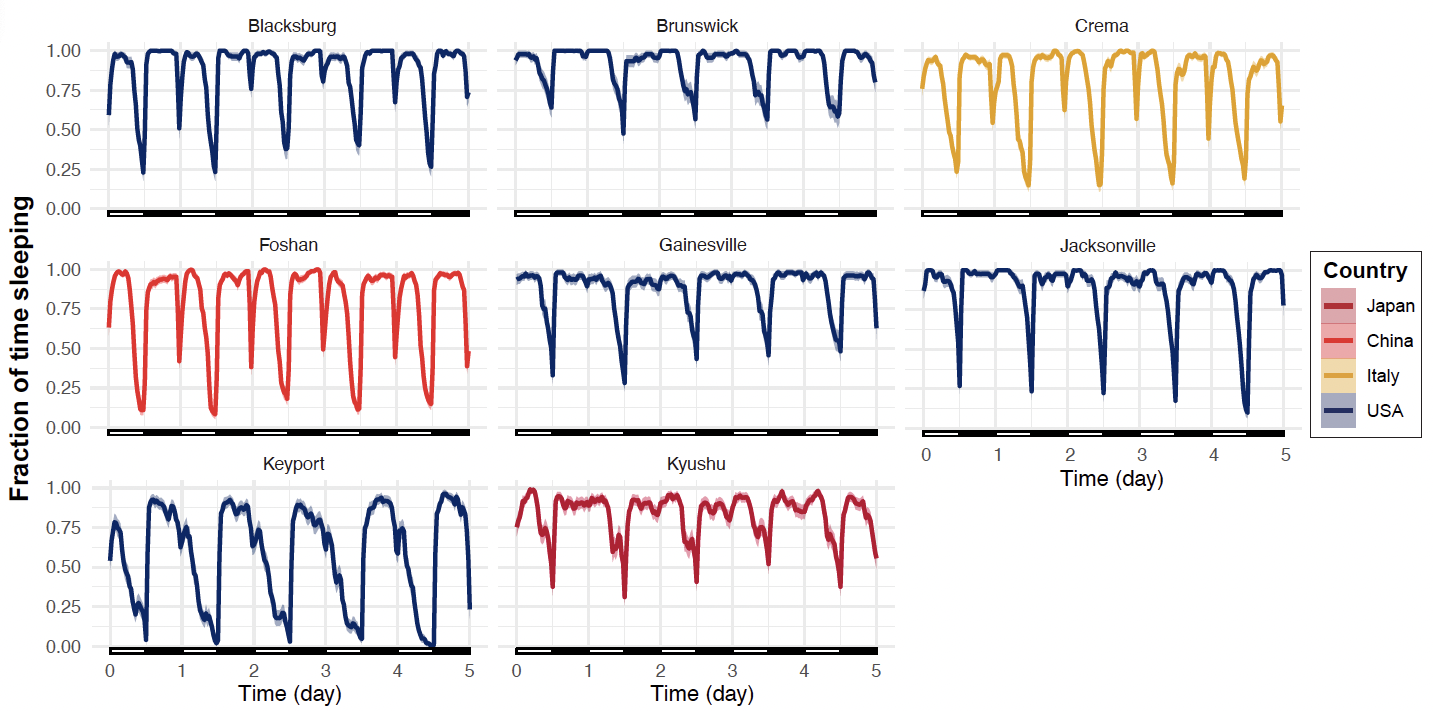

**Figure S2.** Normalized average fraction of time spent in a sleep-like state during the day across five consecutive days of the experiment. The white rectangles below the chart indicate the *Zeitgeber* times when the light is ON, and the black rectangles indicate times when the light is OFF. The shaded area around the curve represents the 95% confidence interval. Each line is color-coded as a function of the country of origin of the population.

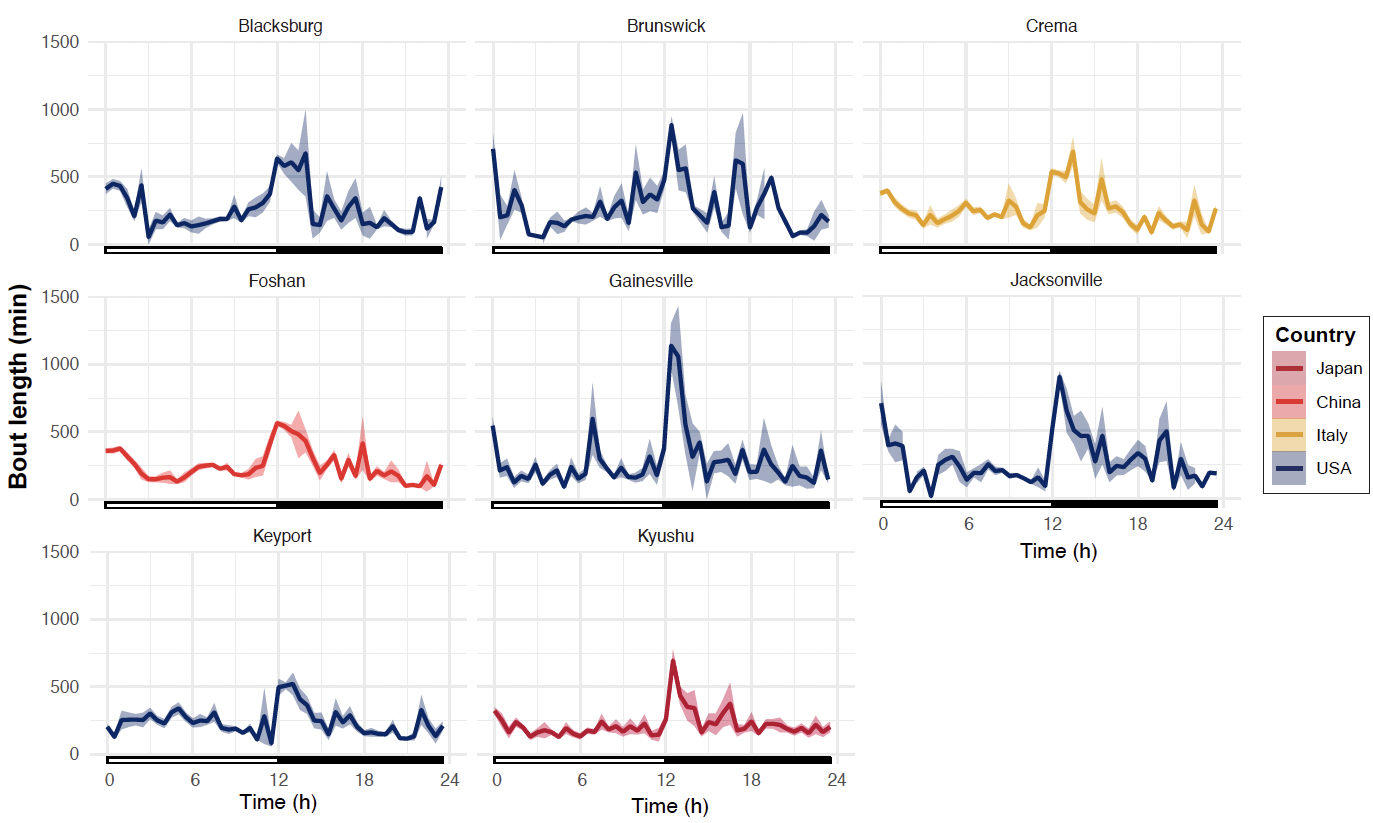

**Figure S3.** Average sleep bout duration during the day. The white rectangles below the chart indicate the *Zeitgeber* times when the light is ON, and the black rectangles indicate times when the light is OFF. The shaded area around the curve represents the 95% confidence interval. Each line is color-coded as a function of the country of origin of the population.
